# Application of Machine Learning and Virtual Reality for Volumetric Analysis of Arterial Lesions

**DOI:** 10.1101/2022.12.23.521811

**Authors:** Ana E. Cartaya, Sophie Maiocchi, Nicholas E. Buglak, Sarah Torzone, Geri Messinger, Edward S. M. Bahnson

**Affiliations:** Dept. of Pharmacology; Center for Nanotechnology in Drug Delivery; Curriculum of Toxicology and Environmental Medicine; Dept. of Cell Biology and Physiology; University of North Carolina at Chapel Hill, NC 27599, USA

## Abstract

Cardiovascular disease (CVD) remains the leading cause of mortality worldwide. Preclinical studies to research and validate therapeutic interventions for CVD often depend on two-dimensional histological surveys. The use of light sheet fluorescence microscopy together with optical clearing methods amenable to immunofluorescence staining are recent advances, all of which deliver detailed three-dimensional rendering of vessels. This offers the ability to describe and quantify features critical in CVD models, specifically, atherosclerotic plaque burden in atherosclerotic animal models and neointimal hyperplasia in surgical models. The main challenge for this approach remains the lengthy, hands-on, analysis time. Labkit is a user-friendly Fiji plugin that applies a machine-learning algorithm to create 3D renderings from large microscopy data. Likewise, syGlass a virtual reality (VR) software, allows for 3D visualization and analysis of information-rich image datasets. The application of these tools is expected to decrease the hands-on analysis time required to generate accurate volumetric renderings of arterial disease and injury features in animal models of CVD. For atherosclerotic burden analysis, Ldlr^−/−^ (C57/BL6) mice aged 6-8 weeks were fed a high-fat diet for 15 weeks to allow the development of atherosclerotic plaque along the aorta. For neointimal hyperplasia analysis, surgically intervened carotid arteries from rats and mice were collected 2 weeks post-surgery. iDISCO+ or AdipoClear and immunolabeling together with light-sheet fluorescence microscopy allowed for three-dimensional visualization of the vessels. Both Imaris software v9.9.1 and the built-in bridge to ImageJ/Labkit were used to quantify plaque burden and neointimal hyperplasia manually or automatically. syGlass was also utilized for the quantification of plaque burden and other disease-associated characteristics. Our findings indicate that both Labkit and syGlass offer effective and user-friendly platforms for the segmentation of atherosclerotic plaque and/or neointimal hyperplasia in animal models.

## 1. INTRODUCTION

Cardiovascular disease (CVD) persists as the leading cause of mortality in the world, with approximately one-third of deaths worldwide attributed to a cardiovascular event.^1,2^ Atherosclerosis, a progressive inflammatory condition of the arteries, significantly contributes to CVD. The gradual subintimal deposition of proinflammatory lipoproteins and immune cells results in the development of atherosclerotic plaque in the affected arteries.^3^ Thrombosis, a consequence of plaque rupture, can lead to serious clinical manifestations which include heart disease, stroke, and peripheral arterial disease.^3,4^ Severe atherosclerosis often relies on surgical interventions to restore normal blood flow. These approaches fail as a consequence of vessel re-narrowing, termed restenosis, caused by an over-exaggerated healing response, known as neointimal hyperplasia.^5^ Atherosclerosis and restenosis are highly complex, multifactorial diseases, which are difficult to recapitulate with *in vitro* models.^6,7^ As such, animal models are indispensable to understanding the pathophysiology of these cardiovascular diseases and discern effective therapeutic interventions.

Traditionally, methods to quantify atherosclerotic burden rely mainly on histological preparations of aortic root sections, and *en face* analysis of aortas. Similarly, histological methods are used to quantify neointimal hyperplasia in the affected vessels. These approaches are widely and frequently used, however, these methods lack universal standardization; and importantly fail to recapitulate the complex three-dimensional architecture of atherosclerotic plaque or neointimal hyperplasia.^8–10^ Recently, our lab and others validated the feasibility of coupling optical clearing methods amenable to immunofluorescence staining such as iDISCO+,^10,11^ AdipoClear,^12^ and CUBIC^13^ with light sheet fluorescence microscopy (LSFM) for the three-dimensional (3D) rendering and analysis of medium to large-sized murine arteries. Further, our lab reported that LSFM processed arteries can be recovered, rehydrated, cryosectioned, and histologically processed with no significant effect on arterial size.^10^ Also, we showed the feasibility of using the recovered arteries for additional immunofluorescence staining allowing for secondary multiplexing using the same samples. Altogether, LSFM appears the obvious choice for the 3D *ex vivo* analysis of vascular features in animal models of CVD. Recent advances in optical clearing and multi-color immunostaining methods for LSFM have opened the door for scientific discovery in many research areas, ^14–16^ in the process generating vast amounts of data-rich image information, often reaching hundreds of gigabytes in size.^17^ The visualization, handling, and laborious, time-consuming, manual segmentation required for the volumetric analysis of these large and complex datasets remain a major limitation for researchers looking to transition to LSFM.

Conventional visualization tools struggle to accurately depict the intricate multidimensional characteristics of image datasets, often presenting 3D acquisitions on a per-slice basis or as 2D projections. This limitation can hinder the clear interpretation of complex 3D structures and may result in the potential loss of valuable information. To address such difficulties new technologies have emerged to facilitate visualization and analysis. One such new and promising technique, syGlass (www.syglass.io), is a software package that relies on virtual reality (VR) using head-mounted displays.^18^ This tool allows researchers to take advantage of binocular vision to facilitate the handling of large multidimensional image datasets through an intuitive interface, with accessible and affordable hardware.

Additionally, artificial intelligence (AI) methods have become more accessible in recent years across all disciplines.^19^ The latest emergence of powerful microscopic imaging techniques allows for the visualization of biological structures in extraordinary detail, leaving scientists and physicians to decipher the best approach to extract relevant quantitative data from these datasets. To bridge the gap, several AI machine learning based automatic pixel classification and segmentation tools have been developed including CellProfiler,^20^ Ilastik,^21^ QuPath,^22^ Napari,^23^ and Trainable Weka Segmentation^24^. Labkit,^25^ the newest addition to this set of tools, is available as a Fiji plugin and unlike its predecessors, is capable of handling large 3D data without the need for specialized personal computers (PCs). Additionally, Labkit features a user-friendly interface with simple drawing tools hence making the application widely accessible to biomedical researchers.

In this study, we aimed to validate different segmentation approaches to analyze 3D image datasets of different models of vascular injury by comparing manual segmentation in 2D, VR-assisted manual segmentation in 3D, and Labkit’s machine learning algorithm. We show that Labkit derived pixel classification and automatic segmentation is an accurate and reliable method of analysis, henceforth widening the bottleneck in this data-rich approach. Furthermore, both Labkit and VR segmentation delivered segmentation accuracy comparable to manual approaches, while being considerably faster and, in the case of Labkit, amenable to automation. The overall aim of this article is then to further establish a methodology for analyzing common vascular features in animal models of CVD in a timely manner.

## 2. MATERIAL AND METHODS

### 2.1 Materials

Paraformaldehyde (PFA) (O4042-500; Thermo Fisher Scientific). Sodium azide (NaN_3_) (S2002; Sigma-Aldrich). Sucrose (S0389; Sigma-Aldrich). Methanol (MX0475-1; Supelco). Phosphate buffered saline (PBS) (20-134; Apex Bioresearch Products). Glycine (A13816.0E; Thermo Fisher Scientific). Tween-20 (P1379; Sigma-Aldrich). Triton X-100 (X100; Sigma-Aldrich). Heparin sodium (H3393; Sigma-Aldrich). Dichloromethane (DCM) (650463; Sigma-Aldrich). Tris base (BP152; Thermo Fisher Scientific). Acetic Acid (ARK2183; Sigma-Aldrich). Agarose (20- 102GP; Apex Bioresearch Products). EDTA (BDH9232, VWR). Dibenzyl ether (DBE) (148400025; Thermo Fisher Scientific).

### 2.2 Animals

All animal protocols here described have been approved by the Institutional Animal Care and Use Committee (IACUC) of the University of North Carolina at Chapel Hill IACUC-ID (21-147, 18-132 and 16-255). 6-7-week-old male Ldlr^−/−^ mice (B6.129S7-Ldlrtm1Her/J, stock number: 002207), were obtained from The Jackson laboratory. Mice were allowed free access to food and water. After a week of acclimation, Ldlr^−/−^ mice were started on a western high-fat diet consisting of 40% fat, 17% protein, 43% carbohydrate by kcal and 0.15% cholesterol by weight (RD Western Diet, catalog number: D12079Bi). Mice were euthanized, and hearts and aortas were collected at 23 weeks.

### 2.3 Mouse carotid ligation

Surgery was performed as described in ^10^. C57BL/6 mice (stock number: 000664) were obtained from The Jackson laboratory. Mice aged 8-10 weeks were anesthetized using isoflurane (1.5-2.5%). After sterile field preparation, a midline incision was made in the ventral surface of the neck. The left carotid artery just proximal to the bifurcation was exposed, and tightly tied with polypropylene surgical suture (7-0) (XS-P718R11, ADSurgical). The animals received Carprofen for pain management before, and twice a day for 48h after the surgery. Euthanasia occurred 2 weeks post-operative under isoflurane overdose.

### 2.4 Mouse aorta and aortic root collection

To harvest the thoracic aorta, adipose tissue was carefully removed first, then the artery was cut at the aortic root and a few millimeters into the large three branches (brachiocephalic, left common carotid, and left subclavian arteries) while the aorta was excised at the bottom along the line of the diaphragm. Carotids, aortas and hearts were maintained in 4% PFA in 1X PBS at 4°C overnight, followed by two washes of PBS at room temperature the next day.

### 2.5 Rat carotid artery pressure-controlled segmental balloon injury

We performed secondary analysis of rat carotid arteries using the raw data originally acquired by Buglak et al^10^. A detailed visual guide of the balloon injury model is found in ^26^. The details of the experimental procedures, animal surgery and sample preparation can be found in the original articles.

### 2.6 Mouse aorta immunostaining and processing for LSFM

Aortas were processed following the AdipoClear protocol,^27^ with mild modifications. All steps were done at room temperature while shaking in 5 mL microtubes with 5 mL of liquid; except for immunostaining, where samples are incubated in 1.7 mL microtubes with 1.5 mL of diluted antibody. Samples were dehydrated in an increasing methanol (MeOH) gradient (20%→40%→60%→80%→(2x)100%) v/v in B1n buffer (0.3M glycine, 0.1% Triton X-100, and 0.01% NaN_3_ in dH_2_O, pH 7) for 30 minutes per dilution. Samples were delipidated in three 100% dichloromethane (DCM) washes for 1 hour, overnight, and 1 hour the next day. Two washes of 100% MeOH were used to remove DCM, each 30 minutes. For immunostaining, aortas were rehydrated in a decreasing MeOH gradient (80%→60%→40%→20%) in B1n buffer, with two final washes in 100% B1n buffer for 30 minutes per dilution. The rehydrated samples were washed in PTxwH buffer (0.1% Triton X-100, 0.05% Tween 20, 0.01% NaN_3_, and 2 µg/mL of heparin sulfate in 1X PBS) for 2 hours. Aortas were then probed with primary rabbit anti-cd31 (0.25 µg/mL, ab28364, Abcam) and rat anti-mouse cd68 (5 µg/mL, MCA1957, Bio-Rad) diluted in PTxwH buffer for 4 days. Before, in between, and after secondary antibody probing, the samples were washed with PTxwH buffer 6 times for 30 minutes and a final time overnight. Aortas were probed sequentially using secondary donkey anti-rabbit IgG AF790 (10 µg/mL, A11374, Thermo Fisher Scientific) and goat anti-rat PE (50x stock concentration not reported, STAR73, Bio-Rad) diluted in PTxwH buffer, each for 5 days. The nucleic acid stain TO-PRO™-3 Iodide (642/661) (200 nM, T3605, Thermo Fisher Scientific) was added together with the final secondary antibody. Next, aortas were washed in PBS twice for 30 minutes, then warmed to 37°C for 1 hour and carefully embedded in 1% agarose (20-102GP; Apex Bioresearch Products) prepared in 1x TAE buffer. Once solidified, the agarose block containing the arteries was removed and dehydrated in an increasing MeOH gradient (20%→40%→60%→80%→(3x)100%) v/v in dH_2_O for 30 minutes per dilution. The blocks were washed thrice with DCM for 1 hour, overnight, and 1 hour the next day. Arteries were cleared in glass scintillation vials using DBE overnight. Samples were stored in DBE in the dark.

### 2.7 LSFM acquisition

Imaging was done at the Microscopy Services Laboratory Core at UNC-Chapel Hill, in a LaVision BioTec Ultramicroscope II (Miltenyi Biotec, Germany) equipped with zoom body optics, an Andor Neo sCMOS camera (Andor Technology, United Kingdom), and an MVPLAPO 2X/0.5 objective (Olympus, Japan) fitted with a 5.7mm working distance corrected dipping cap allowing magnifications between 1.3-12.6X. Blocks were mounted in a resin sample holder and submerged into a container filled with DBE. Aortas were imaged at 1.3X (0.63X zoom) and illuminated with a single sided, three light sheet configuration, with the waist of the sheet in the center of the aortic arch, an NA of 0.024 (beam waist at horizontal focus = 11.5 µm), and a light sheet width of 100% for even illumination in the y-axis. Images were acquired as two tiles with up to 40% overlap, and with a step size of 5 µm. Four channels were imaged as follows: tissue autofluorescence with a 488 nm laser excitation and a Chroma ET525/50m emission filter; CD68-PE with a 561 nm laser excitation and a Chroma ET600/50m emission filter; TO-PRO™-3 Iodide (642/661) with a 647 nm laser excitation and a Chroma ET690/50m emission filter; and CD31-AF790 with a 785 nm laser excitation and an ET800LP emission filter. Carotids were imaged at 1.6X mag (0.8X zoom) and illuminated with a single sided, three light sheet configuration, with the waist of the sheet in the center of the field of view, an NA of 0.043 (beam waist at horizontal focus = 17 µm), and a light sheet width of 100%. Step size was set at 5 µm. Two channels were imaged as follows: tissue autofluorescence with a 488 nm laser excitation and a Chroma ET525/50m emission filter; and CD31-AF647 with a 647 nm laser excitation and a Chroma ET690/50m emission filter.

### 2.8 LSFM data processing

Files were exported as 16-bit ome.tif stacks and converted to Imaris (Bitplane, Oxford Instruments) .ims files with the Imaris File Converter software (v9.9.1). The Imaris tile files obtained were stitched with Imaris Stitcher (v9.9.1) with the final alignment done automatically or manually as needed. Imaris (v9.9.1 or 10.0.0) was used to visualize three dimensional projections with 3D viewer.

### 2.9 Manual segmentation

Manual plaque volume was determined using the manual drawing tools under the surface creation option in Imaris (v9.9.1 or 10.0.0) (Bitplane, Oxford Instruments). Plaque and neointimal hyperplasia were delimited by cd31 staining in the lumen and diffuse autofluorescence around the border of the inner elastic lamina. As illustrated in Figure 1, internal elastic lamina (IEL), and atherosclerotic plaque contours were drawn along the anatomical transverse and sagittal planes using a combination of the isoline, time or distance drawing mode in Imaris. Surfaces obtained from the transverse and sagittal contours drawn were merged into a single surface with the unify option. Manual VR-assisted 3D segmentation was done using syGlass (v1.7.2) (IstoVisio Inc, Morgantown, WV). Plaque segmentation was performed using the autofluorescence channel (Figure 1). In the autofluorescence channel in maximal projection intensity (MIP) mode, brightness, contrast, window, and threshold were adjusted to visualize only the bright elastic lamina. The media was masked using the cut plane tool. Masking was reviewed and curated in 3D view. Using both the autofluorescence channels in MIP mode, brightness, contrast, window, and threshold were adjusted to visualize the plaque and not the media. The plaque was masked using the cut plane tool. The mask was reviewed and curated in 3D view. Mask statistics were exported to obtain quantitative measures of plaque volume. Surfaces or meshes were created from the masks for visualization and further analysis.

**Figure 1.**
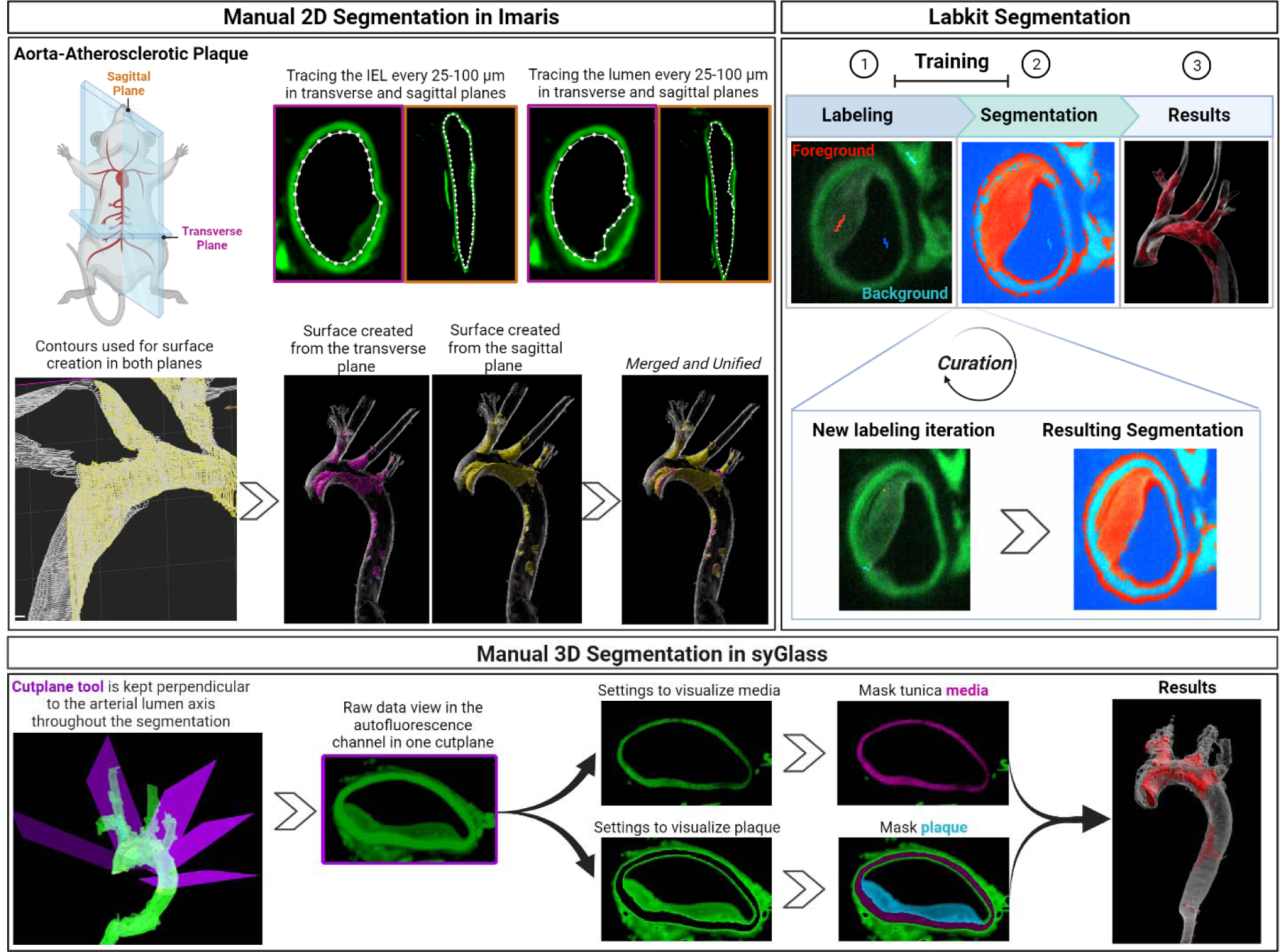
Overview of the image segmentation process. Manual 2D Segmentation in Imaris (Top Left) Atherosclerotic plaque volume was obtained from the merged and unified sagittal and transverse plaque surfaces created with the contours option in Imaris. Labkit Segmentation (Top Right) Step 1. Scribble labeling for binary classification of the plaque or neointimal hyperplasia (foreground, red) and everything else (background, blue) results in automatic segmentation along the artery as shown in the next step. Step 2. Orange labeling indicates plaque volume, whilst blue labeling indicates tissue that is not plaque as determined by the algorithm after training. Steps 1 and 2 are repeated for more accuracy (Curation) Step 3. Labkit automatically segments the plaque volume in the 3D space of the entire aorta. The final result is the automatic production of a plaque volume surface, labeled here in red. Manual 3D segmentation in virtual reality using syGlass (Bottom) brightness, contrast, window, and threshold are adjusted to visualize and mask only the tunica media or the atherosclerotic plaque. Masking for both media and plaque is facilitiated by the use of the cutplane tool, which is kept perpendicular to the lumen throughout the segmentation process, enhancing compatibility with variable vessel anatomy.

### 2.10 Labkit segmentation

Segmentation for plaque and neointimal hyperplasia was done using the autofluorescence and endothelium (cd31) channels as shown in Figure 1. Macrophage distribution in plaque was determined with a combination of the macrophage (cd68), endothelium (cd31), and autofluorescence channels. Necrotic core or acellular regions were determined using a combination of the nuclei (TO-PRO-3), endothelium (cd31), and autofluorescence channels. For all, binary labeling both foreground voxels (plaque, macrophages, or necrotic core) and background voxels (everything else) was done with a brush size of 1 for precise labeling. As illustrated in the Labkit Automatic Segmentation Guidelines imageJ repository (https://imagej.net/plugins/labkit/guidelines) the algorithm performs poorly with large training data so it is very important to label only a few representative objects with the thinnest lines, avoiding pixels at the object edges which carry the potential to confuse the classification algorithm. Generally, the longer the vessel the more the intensity variability, so the focus should be first on finding and sparse labeling a representative area where there is visible difference between foreground and background, and in pixels intensity gradient for both labels. Default basic filters (original image, gaussian blur, difference of gaussians, gaussian gradient magnitude, laplacian of gaussian, and hessian eigenvalues) and sigma values (1, 2, 4, 8) were used for training, as well as the option for GPU acceleration using clij/clij2 whenever compatible with the in-use GPU. After the initial training, the user should navigate the vessel volume, performing additional sporadic labeling and training (curation) as needed before retrieving the final volume. The Imaris software provides a useful tool to store the Labkit processing parameters from the resulting surface when it is generated in Imaris using the bridge to Labkit (Fiji). To utilize this feature, in the Labkit-derived surface under the Create menu, select the option to store parameters for batch processing. Storing the parameters enables Labkit processing of a region of interest without needing to retrain the model.

### 2.11 Aortic root histological processing, staining, and analysis

After perfusion-fixation and overnight 4% PFA incubation at 4°C as described above, hearts were bluntly cut at an angle parallel to the aortic sinus. The section containing the aortic root was transferred to 30% sucrose for overnight incubation at 4°C. Tissue was frozen in O.C.T. and stored at −80°C until processing. Frozen aortic roots were cut in 10 µm sections, 6 sections over 9 slides serially interrupted as recently recommended by the American Heart Association.^28^ Sectioning was done at Histology Research Core Facility at the University of North Carolina at Chapel Hill. Matching slides per animal were immunofluorescence (IF) stained for cd68 (5 µg/mL, MCA1957, BioRad) followed by donkey anti-rat CF750 (2 µg/mL, 20857, Biotium). Slides were imaged with the Slideview VS200 (Olympus, Japan) at the Hooker Imaging Core at the University of North Carolina at Chapel Hill. Plaque, and macrophage content (cd68+ areas) were measured using FIJI (https://imagej.net/software/fiji/) and normalized against secondary antibody only internal controls.

### 2.12 Statistical analysis

Statistical analysis was conducted in Graphpad Prism version 9.3.1 (Graph Pad Software Inc., San Diego, CA). Analysis of any three groups was done using one-way ANOVA. The best fit line between selected values was determined using a simple linear regression, and the resulting R squared (R^2^), p value and slope (m) reported. Correlation between more than 2 variables was assessed firstly with the multiple correlation coefficient, with post hoc linear regression between individual pairs.

## 3. RESULTS

The aim of this article is to validate Labkit, a user-friendly application of machine learning for automatic pixel classification and image segmentation, against VR-assisted segmentation and manual 2D segmentation for the accurate volumetric analysis of common vascular features gathered from LSFM processed arteries. The Labkit VR, and manual 2D analysis workflow is illustrated in Figure 1, Labkit utilizes both Fiji and the integrated Labkit bridge to the Imaris software available on version 9.0 or higher, whereas the VR analysis was performed in syGlass v1.7.2.

### 3.1 Volumetric analysis of atherosclerotic plaque with machine learning based automatic pixel classification vs their manual counterparts

Firstly, we once again validated the combination of LSFM and AdipoClear clearing protocol for the imaging of multiplexed immunostained thoracic aortas of high diet fed athero-prone mice. LSFM proves an outstanding approach to parse out anatomical details in these samples, as shown by the clear 3D projections in Figure 2A, and the original optical slices of aortic arch and descending aorta (Figure 2B) insets of the aortas processed. The optical slices, in particular, clearly show atherosclerotic plaque as the area delineated by the overlap of diffuse tissue autofluorescence (488 nm channel) and a bright cd31+ endothelium layer.

**Figure 2.**
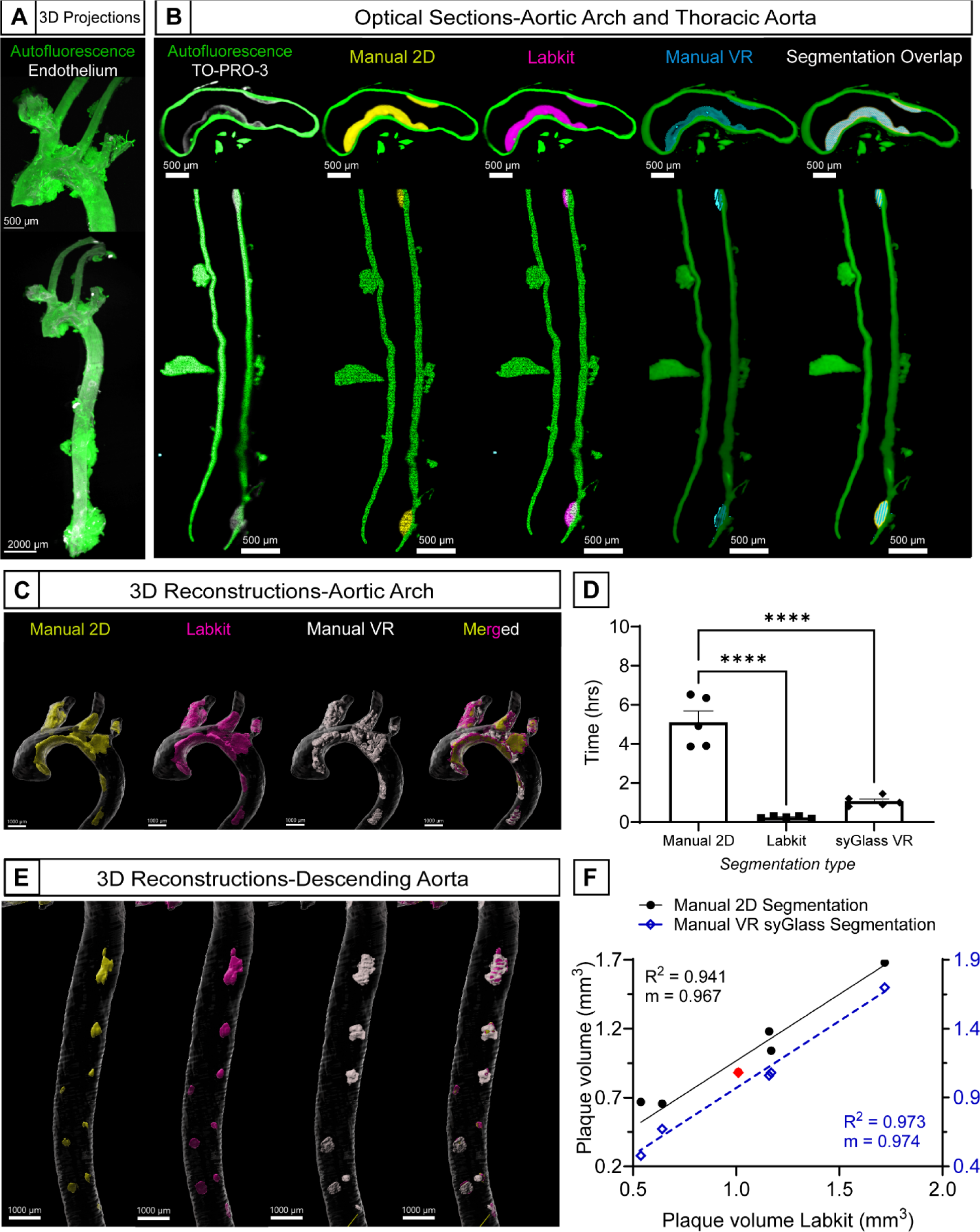
Labkit and syGlass based volumetric analysis results in accurate labeling of atherosclerotic plaque with a significantly shortened hands-on analysis time. (A) 3D volume of a representative mouse aortic arch (top, scale bar = 500 µm) and complete thoracic aorta (bottom, scale bar = 2000 µm) obtained using light sheet fluorescence microscopy. (B) Optical sections representatives from a single aorta (top) aortic arch, and (bottom) thoracic descending aorta showing atherosclerotic plaque delineated by both tissue autofluorescence and the nuclear stain TO-PRO-3, together with manual 2D (Imaris), Labkit or manual 3D (VR, syGlass) based segmentation and their overlap. Scale bar = 500 µm (top and bottom). (C) 3D reconstructions of manual 2D (Imaris), Labkit or manual 3D (VR, syGlass) based segmentation of atherosclerotic plaque in a representative mouse thoracic aorta with a zoomed view into atherosclerotic plaque rich areas of the aortic arch and (E) the descending aorta. Scale bar =1000 µm. (D) Hands-on analysis time taken for manual 2D (Imaris), Labkit or manual 3D (VR, syGlass) segmentation (means ± SEM, n=5, ****p < 0.0001). (F) Best fit line of the linear regression between Labkit and 2D manual segmentation (n=6, R^2^=0.941, m=0.967, ****p < 0.0001) and between Labkit and 3D manual (VR, syGlass) (n=6, R^2^=0.973, m=0.974, ****p < 0.0001) Representative aorta from (B,C,E) shown in red in (E).

To determine the accuracy of Labkit’s output to manual segmentation, aortas were manually labeled in Imaris using the contours function in the anatomical axial and sagittal planes (Figure 1). This was done because we found, on average, a 10% difference in the plaque volume obtained from doing only the axial plane when compared to both planes. Contours from individual planes were transformed into surfaces that were then merged and unified resulting in the final manual segmentation volume. To compare the agreement of Labkit’s output with VR-assisted manual segmentation, aortas were also segmented in syGlass. The cut plane tool was used for plaque visualization and kept at an approximate 90° angle with the axis of the artery using the non-dominant hand of the user while segmentation was performed by masking voxels in the autofluorescence channel using the dominant hand of the user. The masking result was edited and curated in 3D. The 3D corroboration of the segmentation accuracy is an essential step where the power of binocular vision comes into play. For Labkit segmentation the entire thoracic aorta was sparsely labeled, manually classifying a subset of pixels in three dimensions (voxels) as foreground, for areas of plaque, and background for everything else. After the initial training, the curation process, consisting of further labeling and training, was performed sporadically throughout the vessel, supervising the algorithm for proper labeling. Once satisfied with the classifier, a one button operation takes care of automatically segmenting the entirety of the vessel in all dimensions resulting in the Labkit segmentation volume. It is crucial that the resulting Labkit volume be scrutinized for accuracy, as oftentimes edits need to be made prior to retrieving the final volume. Edits in Imaris are simple, using the available surface editing tools. In general, the resulting Labkit surface will include areas outside the arteries as it is the case that any remaining adventitial adipose tissue will have a similar voxel intensity and local contrast characteristics as plaque and be recognized by the algorithm as such. Hence, it is important that excess adipose tissue be removed during the initial dissection of the vessel. The simplest method found to overcome this limitation is to save the trained classifier for batch processing within Imaris and generate smaller surfaces along the aorta excluding as much tissue outside the vessel as possible and lastly, merge all the resulting surfaces. Optical slices of the manual 2D, manual VR 3D, and Labkit segmentation, as well as their overlap, from a representative aortic arch and thoracic descending aorta (Figure 2B) illustrate a high level of agreement between the three methods. Detailed 3D reconstructions of the plaque generated through manual 2D, manual VR 3D, or with Labkit, together with their resulting merged or overlapped surfaces (Figure 2C, 2E) also serve to illustrate this point. Volumetric results of plaque segmentation using the three methods described show a high multiple correlation coefficient of R_labkit, VR3D/Manual2D_= 0.992. Regression analysis of the Labkit measures with 2D Manual and manual 3D VR-assisted segmentation (Figure 2F) reveal a strong linear relationship (n=6, m_manual2D_=0.967, R^2^=0.941, m_VR3D_=0.974 R^2^=0.973, p < 0.0001) Both slopes are not statistically different from 1, validating the accuracy of the algorithm. Moreover, both Labkit and VR-assisted segmentation significantly decrease hands-on analysis time by more than half the time taken to achieve the same manually in 2D (Figure 2D). The ability to perform volumetric analysis of plaque burden enables comparisons that were not readily possible before. We sought to assess the relationship between plaque burden in the aortic arch and plaque in the left subclavian, left common carotid, and brachiocephalic arteries. Brachiocephalic artery (BCA) plaque burden correlated with total plaque in the arch (n=5, m=0.396, R^2^=0.804, p < 0.05) (Sup. Figure 1A, 1B). However, there was no significant correlation between plaque in the arch and the left subclavian (n=5, m=0.038, R^2^=0.495, p=0.184) and left carotid arteries (n=5, m=0.156, R^2^=0.753, p=0.056) (Sup. Figure 1A, 1C). Regardless of the method of segmentation, 3D analysis of atherosclerosis is a new methodology. Hence, we also validated the correlation of 3D analysis with traditional histology of the arotic root. The area under the curve of H&E-stained slides from the aortic root correlated with the volume obtained by either Labkit pixel classification or spyglass VR-assisted segmentation (Sup. Figure 2A, 2B).

### 3.2 Macrophage accumulation and necrotic core distribution in atherosclerotic plaque

An additional major advantage provided by LSFM over traditional *en face* processing is the ability to stain the arteries for multiple targets.^10,11^ Macrophage accumulation and necrotic core distribution in plaque are common queries among atherosclerosis researchers.^29^ We sought then to determine if LSFM coupled with Labkit’s analysis could feasibly provide a volumetric analysis of macrophage accumulation and necrotic core distribution in cd68+, and TO-PRO-3 (nuclei) labeled aortas. Optical sections (Figure 3A) and 3D reconstructions (Figure 3B) of the macrophage (cd68+) layer show extensive distribution of macrophages throughout the volume of atherosclerotic plaque. We found that plaque and macrophage volumes show a strong positive linear relationship (n=6, m=0.452, R^2^=0.971, p < 0.001) (Figure 3C) increasing with plaque volume. We obtained macrophage volumes using VR-assisted segmentation and found a strong linear correlation with the Labkit obtained volumes (n=6, m=1.011, R^2^=0.953, p < 0.001) (Figure 3D). The slope obtained is not different from 1, suggesting that both segmentation methods measure the same variable equally. Additionally, quantification of cd68 immunofluorescence sections in the aortic root, showed a linear correlation between macrophage infiltration in aortic root (volume determined as AUC), and cd68+ volume in the aortic plaque quantified by Lablkit (n=6, R^2^=0.727, m=4.927, p < 0.05) (Sup. Figure 3A). Distribution of acellular regions, characteristic of necrotic cores, appear localized to the aortic arch, where the most severe plaque develops (Figure 3B and 3C). No correlation was found between total necrotic core and plaque volume using either Labkit (Sup. Figure 4A), or syGlass (Sup. Figure 4B).

**Figure 3.**
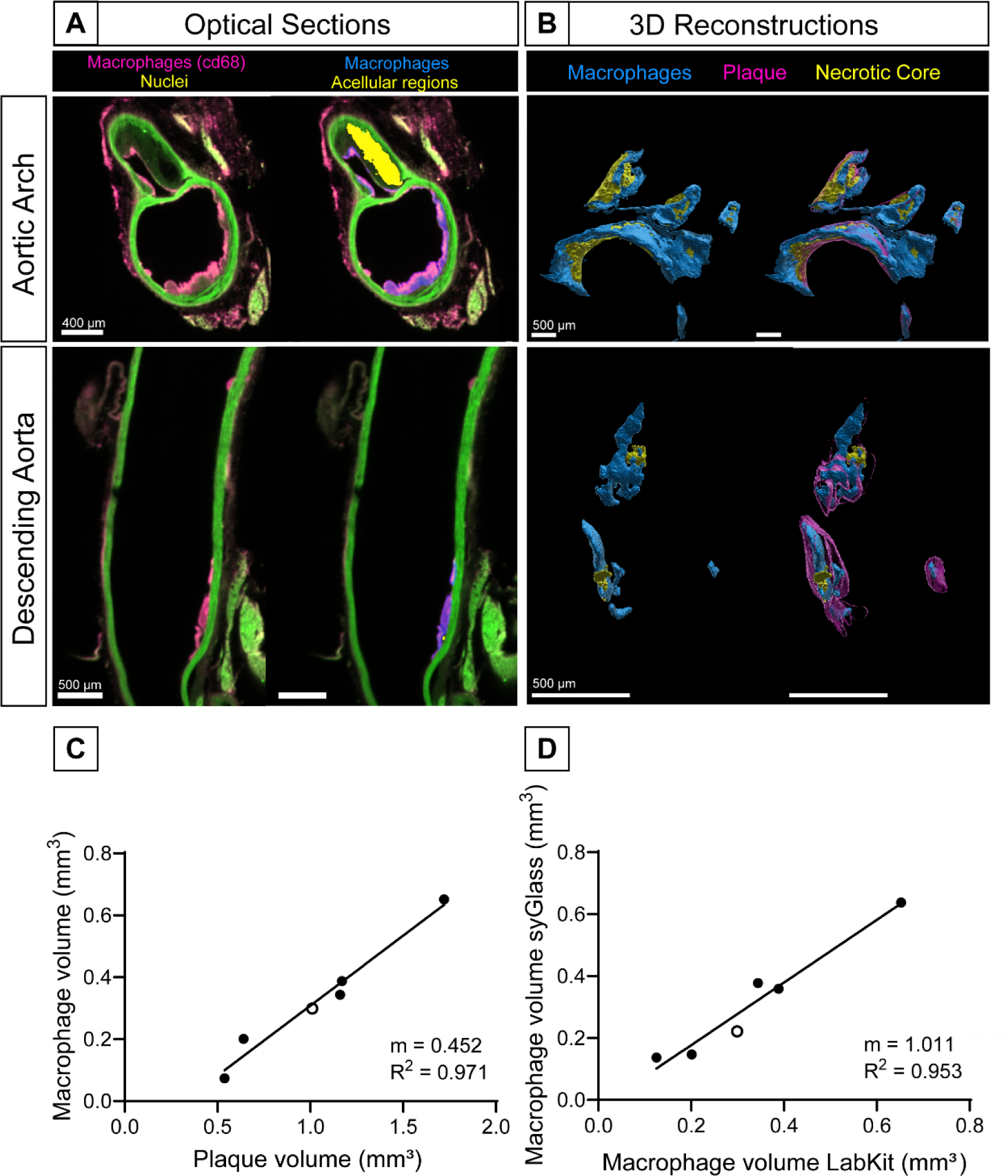
Labkit and syGlass-enabled volumetric analysis of plaque-associated macrophages reveals a linear relationship with plaque volume. (A) Representative optical sections from a single aorta (top) aortic arch, and (bottom) descending aorta displaying macrophage infiltration (cd68), and nuclei (TO-PRO-3) in atherosclerotic plaque delineated by tissue autofluorescence, compared to the same optical section with the resulting Labkit segmentation for macrophages and acellular regions or necrotic core. Scale bar = 400 µm (top) scale bar = 500 µm (bottom) (B) 3D reconstructions of a representative mouse thoracic aorta with Labkit derived plaque, necrotic core, and macrophage volumes in (top) aortic arch, and (bottom) descending aorta, plaque shown as a semitransparent surface. (C) Best fit line of the linear regression between Labkit derived plaque and macrophage volumes (n=6, R^2^=0.971, m=0.452, **p < 0.01). (D) Best fit line of the linear regression between syGlass and Labkit retrieved macrophage volumes (n=6, R^2^=0.953, m=1.011, ***p < 0.001)

### 3.3 Volumetric analysis of neointimal hyperplasia

We also sought to determine the applicability of Labkit’s segmentation to other features of CVD, specifically neointimal hyperplasia in mice and rat carotid arteries. In a previous method from our lab,^10^ we presented a manual method of quantifying both neointimal hyperplasia and medial remodeling consequence of restenosis. This method relied on manually segmenting three different layers of the artery: the lumen (as demarcated by cd31), and the inner and external elastic lamina visible in the autofluorescence channel. Is important to note that because of the concentric nature of neointimal hyperplasia development in the animal models used, it is generally not possible to segment this feature directly. Here, we applied Labkit to neointimal hyperplasia developed in rat carotids (Figure 4A) which shows great correlation with the volume obtained from the reported mask subtraction manual method (Figure 4B) (n=4, R^2^=0.970, m=1.015, p < 0.05). Additionally, we applied the algorithm to ligated mouse carotid arteries 2 weeks post-surgery. In contrast with surgically intervened rat carotids, we found a great degree of medial thickening and remodeling in ligated mouse carotids, leading to loss of voxel contrast between the media and the neointima (Sup. Figure 4). This ultimately led to inaccurate direct segmentation of neointimal hyperplasia. However, a mask subtraction approach using the manual segmentation of the volume of the internal elastic lamina was sufficient to overcome this limitation. We sought then to determine if this approach (Labkit surface-IEL) would be an improvement over previous methods, thus it was directly compared to the native Imaris surface creation tools. Compared to the autofluorescence channel (488 nm) absolute intensity-based surface creation option in Imaris (n=5, R^2^=0.998, m=0.901, p < 0.001), the Labkit derived volume transformation is more accurate (n=5, R^2^=0.999, m=0.978, p < 0.0001) and it is still an improvement over our previously reported methodology in both time and accuracy. This difference is much more striking when comparing the volumes obtained for the media with absolute intensity-based surface creation option in Imaris (n=5, R^2^=0.706, m=0.722, p= ns) which underperforms compared to the Labkit derived volume transformation (n=5, R^2^=0.954, m=1.067, p < 0.05) (Sup. Figure 4).

**Figure 4.**
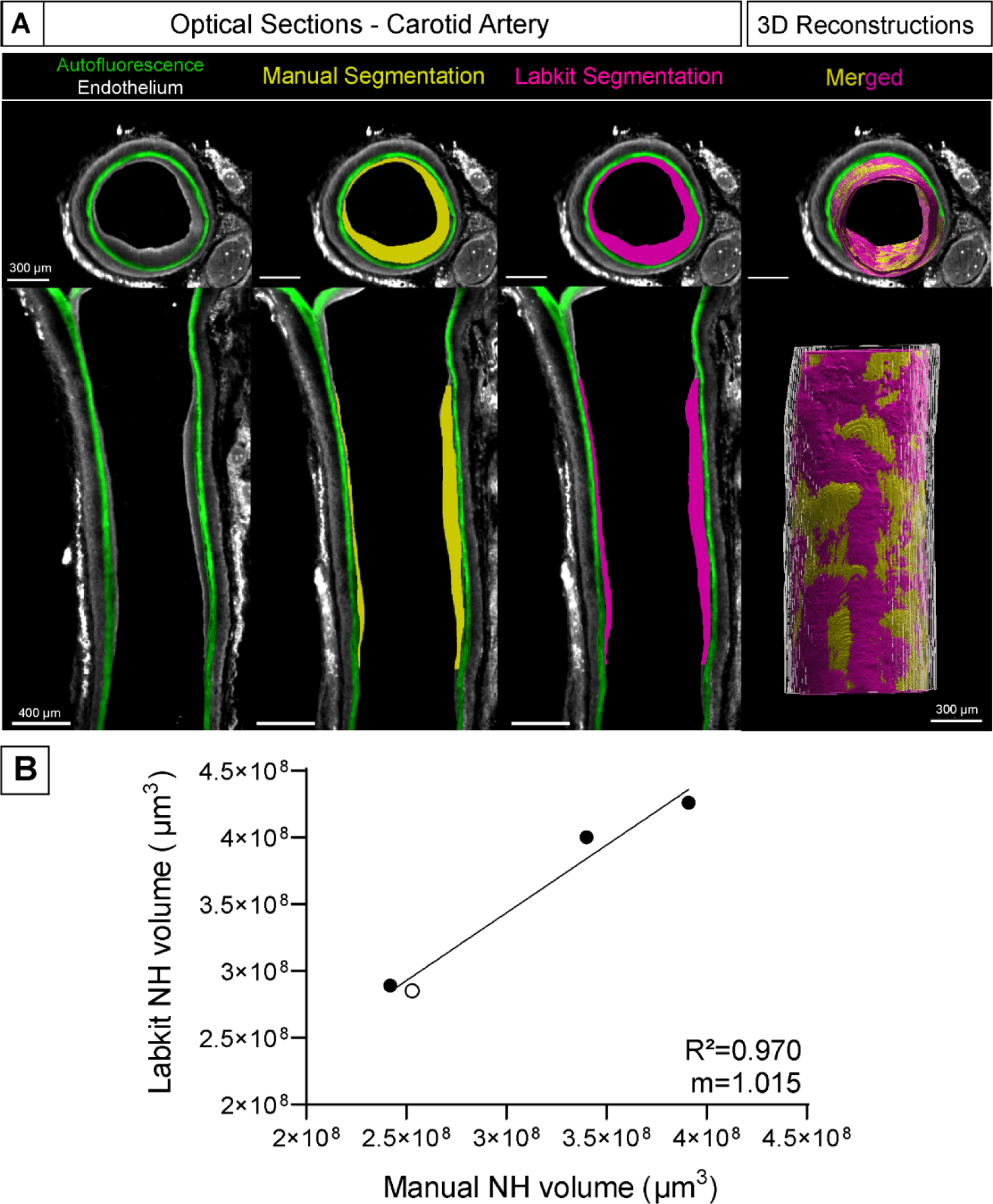
Labkit based volumetric analysis results in accurate labeling of neointimal hyperplasia in a rat model of restenosis. (A) Representative optical sections from a single rat carotid showing neointimal hyperplasia delimited by autofluorescence (green) and cd31 endothelium (white); together with manual (yellow), or Labkit based segmentation (magenta) (top, scale bar = 300 µm) (bottom, scale bar = 400 µm), and a merged 3D reconstruction of the hyperplasia. Scale bar = 300 µm. (B) Best fit line of the linear regression between manual and Labkit derived segmentation (n=4, R^2^=0.970, m=1.015, *p < 0.05).

## 4. DISCUSSION

Herein, we describe the application of a machine learning tool, Labkit, for the quantitative analysis of common vascular features in animal models of CVD, and compare it to both, the traditional manual 2D segmentation technique, and the other cutting-edge tool: VR-assisted segmentation. Specifically, we describe the volumetric analysis of atherosclerotic plaque burden in the aortas of atherosclerotic-prone mice, and neointimal hyperplasia development in the carotids of rat and mice post-surgery. Firstly, we demonstrated that the Labkit derived volumes are significantly correlated with the volumes resulting from manual 2D analysis, and VR-assisted 3D analysis, confirming the accuracy and reliability of Labkit for volumetric analysis of arterial lesions. Importantly, we showed that plaque volumes in the arch and thoracic aorta obtained by Labkit pixel classification, positively correlate with the plaque in the aortic root determined by traditional histological methods in agreement with established results.^30^ Lastly, the implementation of our protocol results in a significant reduction in the time required to analyze plaque volumes, (on average 4 hrs to less than 1 hr), with this including short periods of hands-off training. This further confirms the utility of Labkit derived plaque volume measurements. Likewise, we found that VR-assisted segmentation reduces hands on analysis time compared to manual 2D segmentation, and it also achieves highly correlated results. Finally, we demonstrate that this protocol can be multiplexed for analysis of other important plaque characteristics such as inflammatory cell and necrotic core distribution. Overall, we present two methods that significantly reduce the time to analyze plaque and neointimal hyperplasia volumes from 3D LSFM datasets. Labkit requires neither specialized PCs nor specialized computational skills and is therefore a step toward making LSFM more widely adopted as an accurate volumetric quantitation technique. Similarly, syGlass does not require expensive hardware, needing similar specifications to that of a home gaming PC. Both approaches enable a broader utilization of 3D datasets in CVD research.

Atherosclerosis is a complicated disease, with numerous factors contributing to its development. Unsurprisingly, even the latest established *in vitro* models fail to fully recapitulate the disease.^6^ As such, preclinical animal models, such as the Ldlr^-/-^ and ApoE^-/-^ mice, are essential in aiding our understanding of atherosclerotic disease initiation and progression. Both atherosclerotic plaque burden and volume are two key characteristics that are utilized as a measure of either the influence of a genetic mutation or of a therapeutic intervention upon disease. Current widely-used techniques to analyze plaque volume and burden in atherosclerotic mice rely either on histological sectioning of the aortic sinus^31,32^ as developed originally by Paigen and co-workers in 1987,^33^ or *en face* staining of the aorta with lipid staining dyes such as Sudan IV or Oil red O.^34^ Both techniques have significant drawbacks including the introduction of several sources of variability that decrease the accuracy of plaque burden measurements, thereby increasing the variability of an already highly variable disease model. As an example, it has been reported that a slight tilt of 20° in the sectioning angle can lead to an overestimation of lesion sizes by 15% or more.^34,35^

More recently, advanced 3D imaging techniques have come to the forefront to describe and quantitate atherosclerotic plaque and neointimal hyperplasia volume in preclinical models, including high resolution magnetic resonance imaging (MRI),^36,37^ microscopic computed tomography (microCT),^38,39^ and optical projection tomography (OCT).^40–42^ In line with this growing focus on 3D imaging techniques, us and others have demonstrated the utility of LSFM together with clearing protocols amenable to immunofluorescence staining, such as AdipoClear^12^ and iDISCO+^10^, to provide 3D unbiased quantitative measurements of vascular features.^10–12^ Additionally, previous research has demonstrated that there is a significant correlation between plaque burden obtained by *en face* staining, and volume obtained through microCT imaging;^43^ which indicates that 3D techniques could replace *en face* staining as these approaches become increasingly available, user-friendly and lower cost. Thus far, the major limitation of using LSFM for the quantification of plaque burden and neointimal hyperplasia is the large investment on the hand-on analysis time, which limits its application to typically large datasets seen in preclinical studies. The methodology described herein is a step towards reliable quantitation of 3D datasets using novel technologies like an accessible machine learning application and VR-based analysis, both with significantly lower hands-on analysis time.

Previous authors have applied artificial intelligence approaches in both the clinical,^44–46^ and preclinical settings to detect, quantify and classify atherosclerotic plaque. Jurtz and co-authors recently detailed a deep learning model that was trained for plaque quantification in different anatomical segments of the mouse aorta.^11^ This method resulted in reliable and accurate measurements of plaque volume. However, the major limitations of this approach are often the need for expensive, powerful computers and skills from researchers, as well as the need for large training datasets. The field does continue to take incremental steps toward making deep learning models more accessible, and cost-effective.^47^ An alternative machine learning algorithm, random forests, requires much less training data. As such several tools have been developed for the imaging community, all unable to process large image data. Labkit, also an application of the random forests algorithm, fills that gap.^25^ The application features a simple layout for pixel classification and is easily accessed through Fiji to quickly segment large image data. It also features a dedicated documentation page in the ImageJ Wiki (https://imagej.net/plugins/labkit/). Another element result of the shallow learning approach is the passive accumulation of large amounts of supervised ground truth annotations, which in the future can be used for training deep learning algorithms for faster, higher quality output.^25^ Altogether, highlighting the utility of Labkit for the quantitative analysis of atherosclerotic plaque and neointimal hyperplasia.

In addition to atherosclerotic plaque volume and burden, other aspects of atherosclerosis are of relevance. The inflammatory characteristics, including inflammatory cell distribution, as well as necrotic core size, and endothelial erosion are all of interest to quantify. LSFM allows us to multiplex with different molecular markers. Herein we have shown the presence of cd31, cd68 as well as nuclei to probe some of the mentioned characteristics. In particular, we show that macrophage content is highly correlated with plaque volume, an effect previously reported in human atherosclerotic arterial samples.^48^ Lastly, distribution of macrophages appears restricted to the periphery of larger plaques that develop in the aortic arch, in contrast to the smaller plaque that develops in the thoracic aorta which appear more thoroughly infiltrated by the inflammatory cells. These results were observed both using Labkit pixel classification, and VR visualization. Similar observations were made by Jurtz et al, with subtle differences in distribution of cd45 signal.^11^

Important limitations remain for the use of LSFM, Labkit, and syGlass for the volumetric analysis of CVD features. Firstly, current commercially available LSFM equipment, in general, achieves modest cellular resolution compared to well-established techniques like laser scanning confocal microscopy.^49^ LSFM techniques continue to improve at an accelerated rate, with researchers looking to answer questions in subcellular resolution. Recent research suggests that the technology to answer these questions is around the corner, for example, the combination of LSFM and the super resolution technique, structured illumination microscopy (SIM), pushes the lateral resolution to less than 100 nm.^50,51^ Another limitation to the broad adoption of LSFM is the necessary optical clearing step, and validation of compatible antibodies for immunostaining, both of which remain a large time investment.^52^ Additionally, with regards to the Labkit analysis, the application segmentation algorithm classifies pixels independent of their position in the image.^25^ In the case where significant adventitial fat is still present, for example, Labkit may include this tissue in the final plaque volume. As such, it is crucial that the resulting volume be edited for accurate plaque representation, which can add up to an hour to the process. It is possible to overcome this limitation by processing the samples in smaller sections, ultimately merging all resulting surfaces. On the other hand, VR-based analysis using syGlass also presents limitations. The software is relatively new and the only one of its kind. Compared to more established imaging analysis software such as Imaris, syGlass lags behind in number of functions and user-created plug-ins. Additionally, syGlass currently does not support floating point TIFs. This obstacle can be overcome by converting 32 bit floating point TIF stack into 16 bit linear using shareware like FIJI.

In conclusion, we describe and validate the application of a user-friendly machine learning algorithm for automatic 3D segmentation, Labkit, to provide accurate volumetric quantitation of atherosclerotic plaque and neointimal hyperplasia from murine preclinical models of vascular disease. We also describe the use of VR-assisted analysis of 3D datasets to quantify atherosclerotic plaque, acellular necrotic core, and macrophage content. Both approaches outperform manual segmentation. However, Labkit offers automation which could help reduce user bias. Overall, the protocol presented reduces the plaque volume quantitation time to less than an hour. Our technique also allows for the simultaneous analysis of other important plaque features, such as inflammatory cell and necrotic core content by staining for macrophage markers and nuclei, respectively. This greatly increases the practicality of LSFM for the analysis of large data sets of rats and mice, which are typical in studies of atherosclerosis and neointimal hyperplasia development, where the variability typically necessitates 10-15 animals per treatment group. Moreover, Labkit is a widely available Fiji plugin, with no need for specialized PCs, computational skills, or access to large training data sets necessary for deep learning. Thus, literature reports of this application are expected to increase both the appeal of LSFM and these analysis techniques. It is likely this protocol can be successfully applied to other vascular tissue in preclinical vascular disease animal models, including the aortic root. Finally, if need be, in our previous studies we have shown that vascular tissue samples can be recovered, rehydrated, embedded in O.C.T, and sectioned for traditional histological or immunofluorescence analysis, following LSFM imaging.^10^

## ADDITIONAL INFORMATION

The authors declare no conflicts of interest regarding the publication of this manuscript.

## ACKNOWLEDGMENTS

This work was supported by the National Institutes of Health, National Heart Lung, and Blood Institute [Grant K01HL145354]. A.E.C is supported by the National Heart, Lung, and Blood Institute Predoctoral Fellowship [F31HL156427]. S.M is supported by supported by the National Heart, Lung, and Blood Institute Pathway to Independence Award (K99/R00) [K99HL157690]. LSFM imaging was done at the UNC Microscopy Services Laboratory Core supported, in part, by the P30 CA016086 Cancer Center Core Support Grant to the UNC Lineberger Comprehensive Cancer Center, and the North Carolina Biotech Center Institutional Support Grant 2016-IDG-1016. Slide imaging was performed at UNC Hooker Imaging Core Facility, supported in part by P30 CA016086 Cancer Center Core Support Grant. Sectioning and histological processing were done at Histology Research Core Facility at the University of North Carolina at Chapel Hill. The authors thank Mr. Nathan Spencer from syGlass for his assistance with the software and the inclusion of new functionalities needed for our analyses.

## AUTHOR CONTRIBUTIONS

A.E.C, S.M, and E.M.B conceived the study and designed the experiments. A.E.C prepared the first draft of the manuscript. A.E.C, and S.M, collected, stained, and cleared tissue. A.E.C performed imaging, and image quantification analysis. N.E.B performed mouse and rat surgeries, tissue collection, imaging, and manual segmentation. E.M.B. performed imaging analysis, edited the manuscript, and provided the funding. S.T and G.M quantified aortic root staining.

## SUPPLEMENTARY FIGURES

**Supplemental figure 1.**
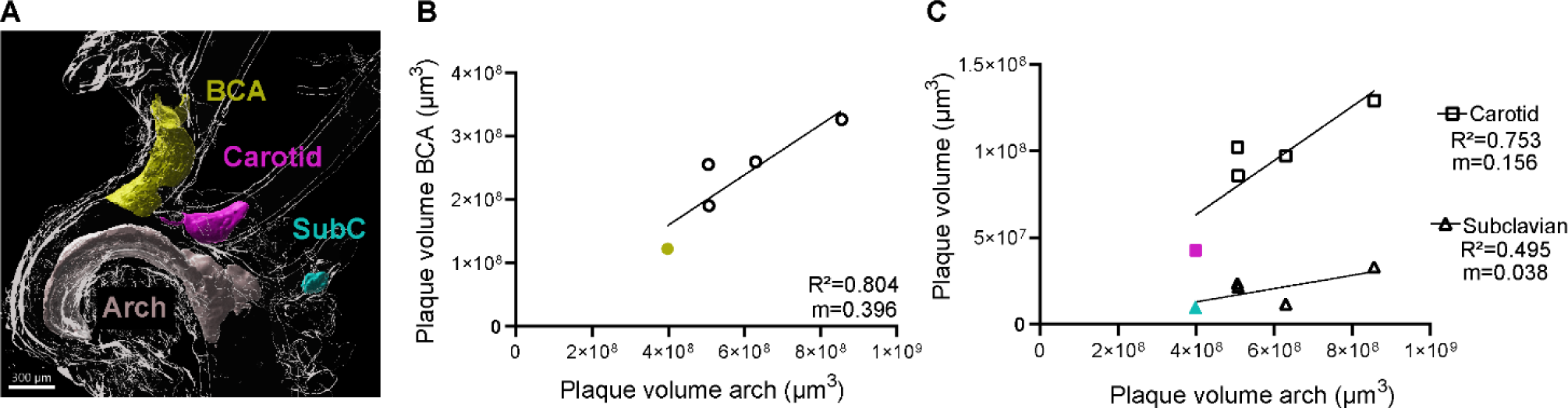
Labkit volumetric analysis describes the atherosclerotic plaque distribution in the aortic arch and associated main arterial branches. (A) 3D reconstruction of the atherosclerotic plaque which develops along the aortic arch (grey), brachiocephalic artery (BCA, yellow), left carotid artery (magenta), and left subclavian artery (cyan). (B) Best fit line of the linear regression between plaque volume in the arch, and the BCA (n=5, m=0.396, R^2^=0.804, *p<0.05) (C) Best fit line of the linear regression between plaque volume in the arch, and the left carotid artery (n=5, m=0.156, R^2^=0.753, p=0.056); or left subclavian artery (n=5, m=0.038, R^2^=0.495, p=0.184).

**Supplemental figure 2.**
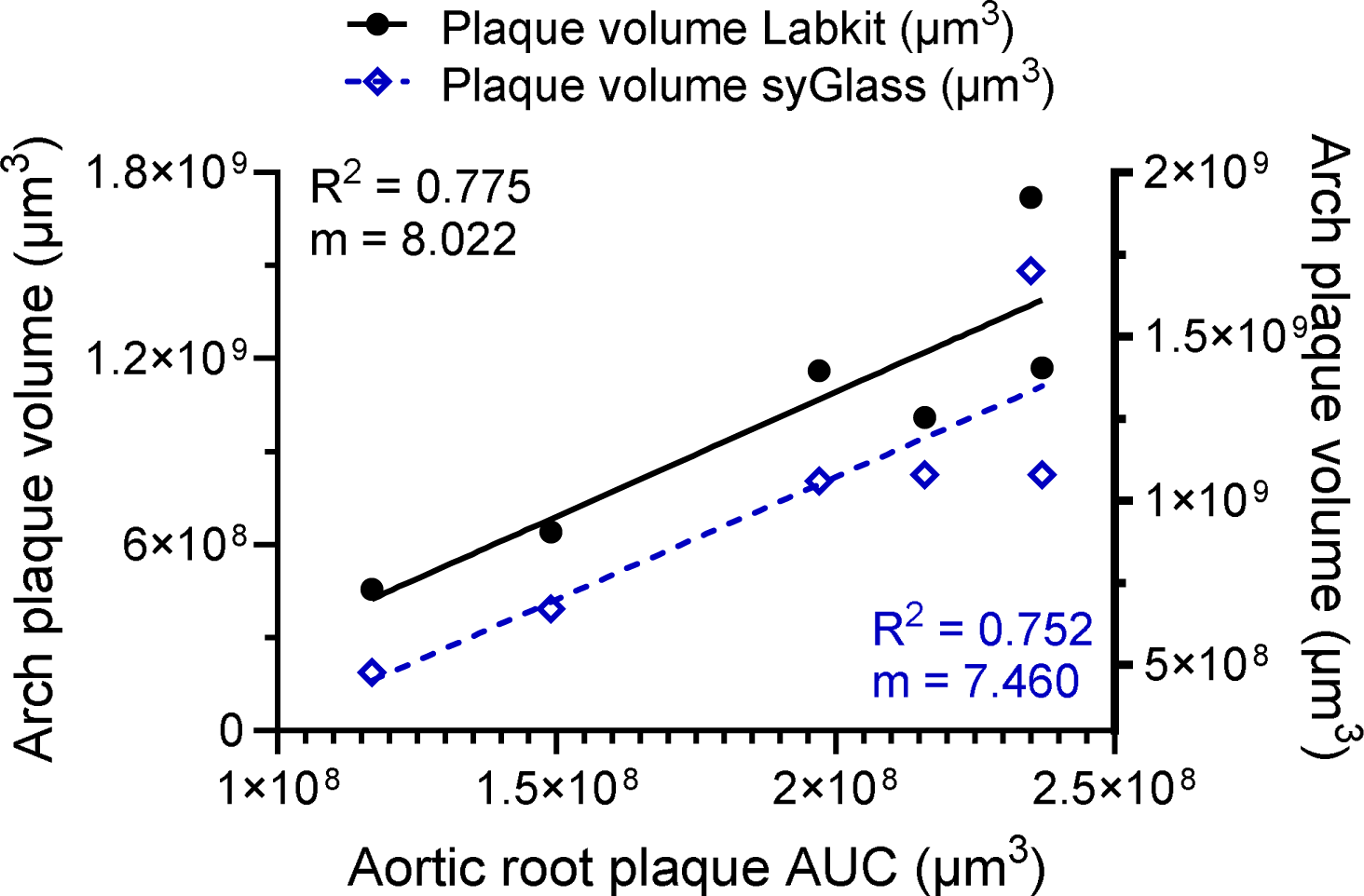
Plaque in the aortic root correlates with aortic plaque volume. Best fit line of the linear regression between area under the curve of aortic root plaque, and aortic arch plus thoracic aorta plaque volume segmented with Labkit. (n=6, R^2^=0.775, m=8.02, *p<0.05) or spyglass (n=6, R^2^=0.752, m=7.46, *p<0.05)

**Supplemental figure 3.**
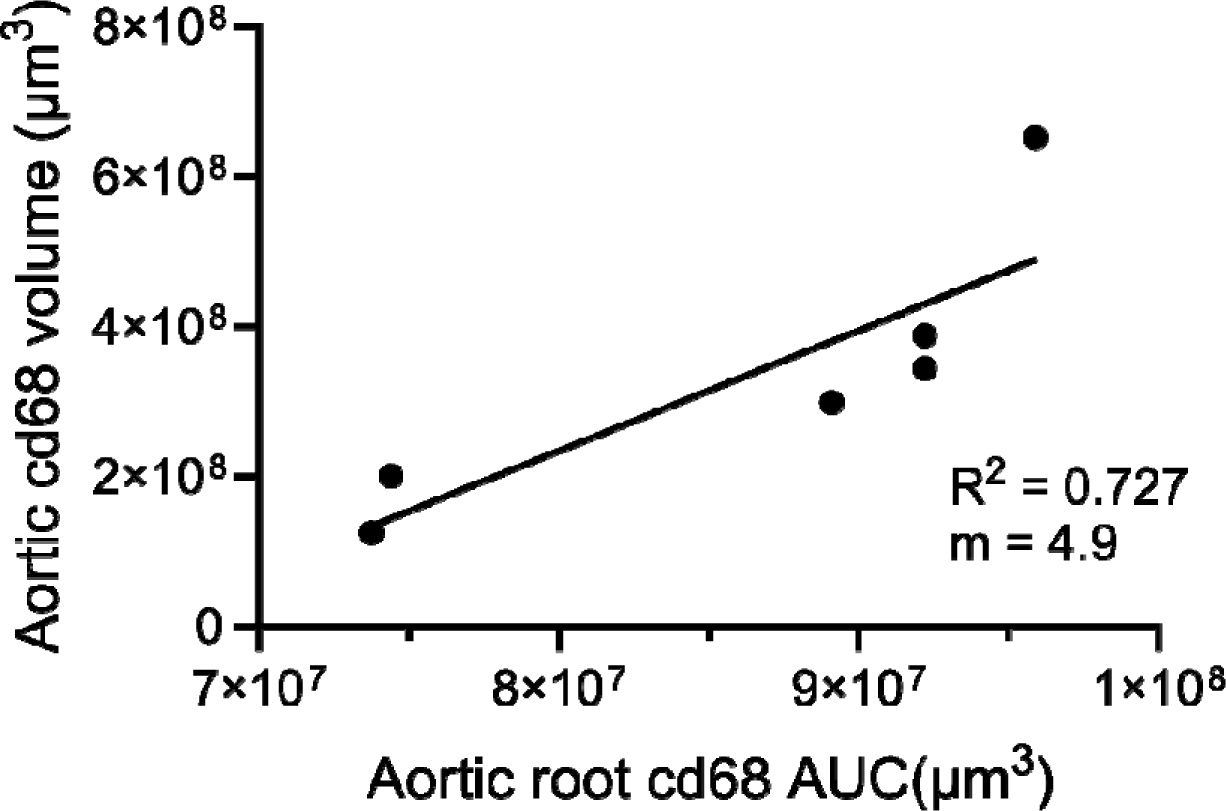
Macrophages in the aortic root correlate with macrophage volume in the aortic arch plus thoracic aorta. Best linear fit between the area under the curve of cd68 positive areas determined by IF in the aortic root and cd68 positive volume in the thoracic aorta and arch determined by Labkit.

**Supplemental figure 4.**
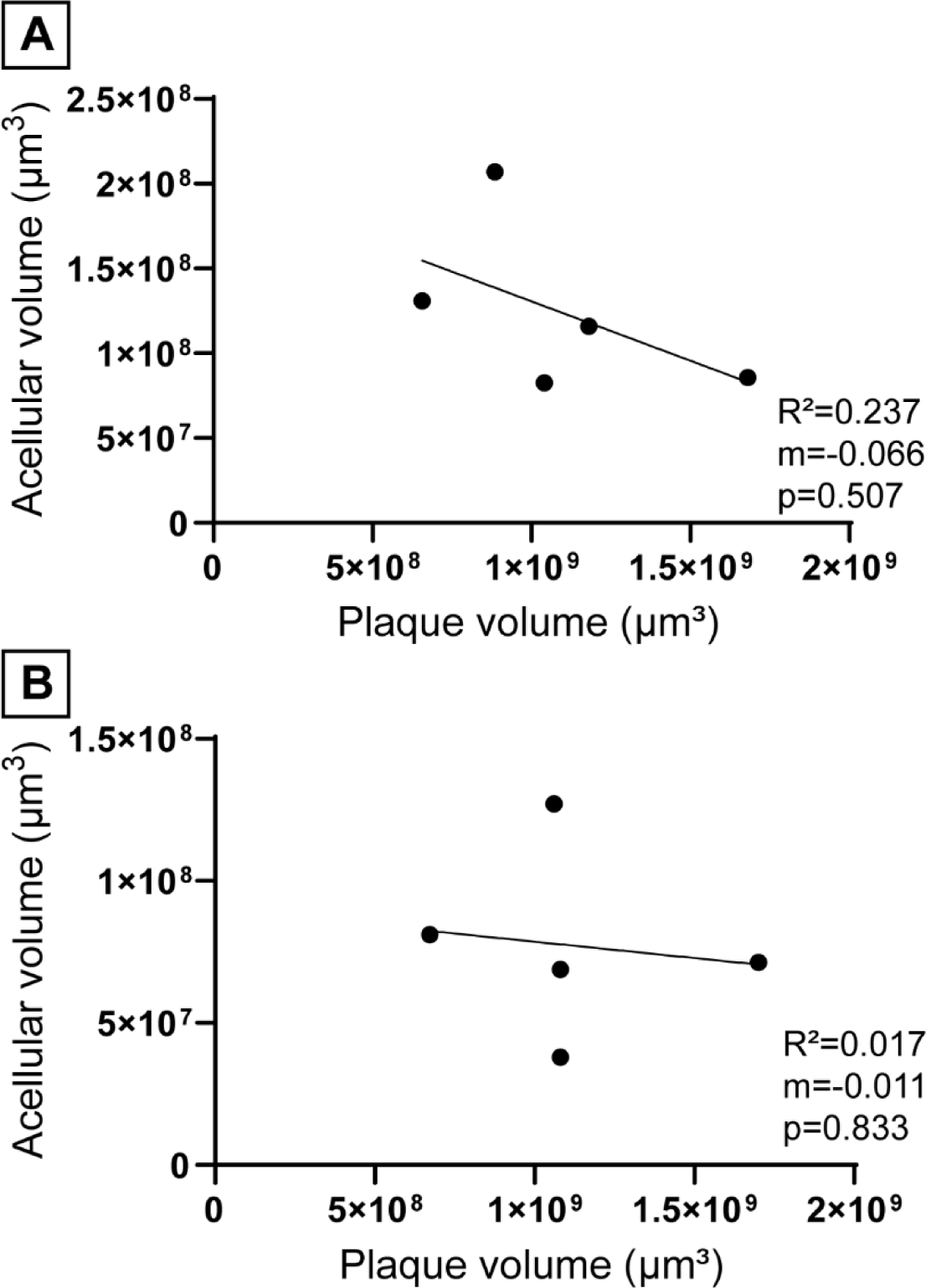
Acellular region volumes do not correlate with atherosclerotic plaque. (A) Best fit line of the linear regression between acellular and plaque volume obtained using Labkit, (n=5, R^2^=0.234, m=-0.062, ns), and (B) VR-assisted segmentation in syGlass (n=5, R^2^=0.017, m=-0.011, ns).

**Supplemental Figure 5.**
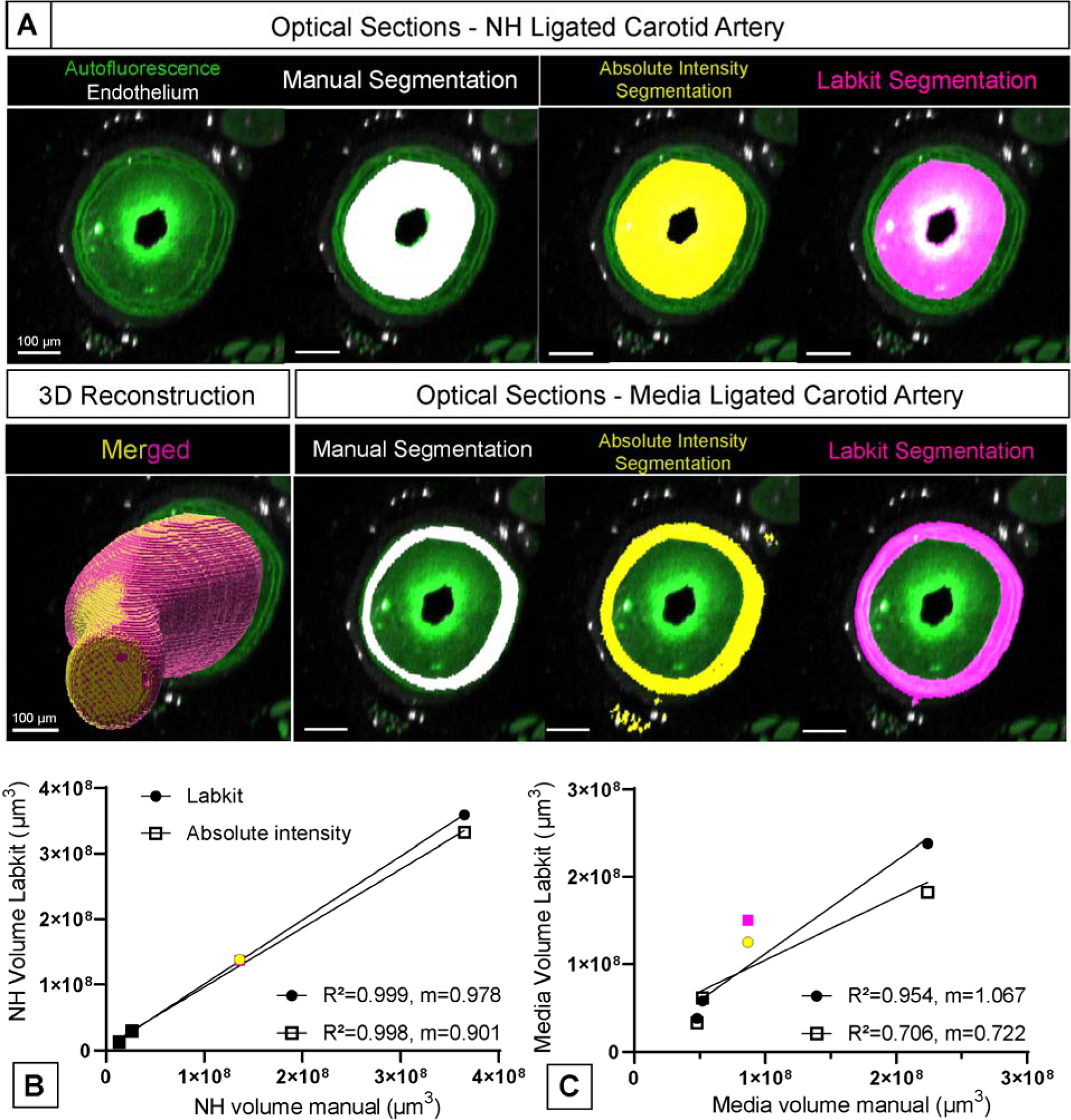
Labkit segmentation coupled with manual segmentation of a single surface produces a more accurate volume of both neointimal hyperplasia and media thickening when compared to Imaris software native tools. (A) Optical sections representatives from a single ligated mouse carotid showing neointimal hyperplasia (top) or media thickening (bottom), together with manual (white), absolute intensity based (yellow) or Labkit based segmentation (magenta), and a merged 3D reconstruction of the hyperplasia. Scale bar = 100 µm. (B) Best fit line of the linear regression between manual absolute intensity based (n=5, R^2^=0.998, m=0.901, ***p < 0.001) or Labkit derived volume (n=5, R^2^=0.999, m=0.978, ****p < 0.0001) of neointimal hyperplasia. (C) Best fit line of the linear regression between manual absolute intensity based (n=5, R^2^=0.706, m=0.722, p=ns) or Labkit derived volume (n=5, R^2^=0.954, m=1.067, *p < 0.05) of the media volume.

